# FRESH FROZEN PLASMA-BASED RESUSCITATION LESSENS LUNG INJURY IN MICE WITH ABDOMINAL SEPSIS AND HEMORRHAGIC SHOCK

**DOI:** 10.64898/2026.07.22.739847

**Authors:** Feng Wu, Jody Cantu, Chavi Rehani, Rosemary A Kozar

**Affiliations:** Shock Trauma Center and Shock Trauma Anesthesiology Research (STAR) Center, University of Maryland School of Medicine, Baltimore, MD

**Keywords:** sepsis, trauma, fresh frozen plasma, neutrophil degranulation, syndecan-1, MMP9

## Abstract

We have previously shown that fresh frozen plasma (FFP) and fibrinogen have protective effects in mice with hemorrhagic shock through restoration of endothelial syndecan-1 and reversal of endothelial injury. In the current study, we tested the hypothesis that a combined model of abdominal sepsis and hemorrhagic shock would induce endothelial syndecan-1 shedding and lung injury which could be attenuated by both FFP and fibrinogen. C57BL/6 mice underwent cecal ligation and puncture (CLP) followed by hemorrhagic shock (HS) and fluid resuscitation with lactated Ringer’s (LR), fibrinogen (5 mg/mouse), and FFP, all at 1X shed blood volume. After 24 hours, lung tissues and plasma were harvested for assays. CLP+HS induced an increase in alveolar thickness and decreases in lung syndecan-1 and lung neutrophil granule-enzymes (myeloperoxidase, neutrophil elastase, and MMP9), with reciprocal elevations in plasma syndecan-1 and plasma neutrophil granule-enzymes (myeloperoxidase, neutrophil elastase, and MMP9). All these alterations were significantly attenuated by FFP but not by fibrinogen. Additionally, CLP+HS-induced hypotension at 24 hours was partially reversed by FFP but not by fibrinogen. FFP administration inhibits CLP+HS-induced neutrophil degranulation to prevent syndecan-1 shedding and lung injury. The current study supports that FFP has therapeutic benefit in a combined septic and hemorrhage shock model.

## Introduction

Sepsis is a life-threatening complication caused by a dysregulated innate immune response to infection, leading to multiorgan dysfunction and death (1). Traumatic hemorrhagic shock is a significant risk factor for sepsis, with over one-third of the survivors developing sepsis (2, 3). Trauma-associated sepsis is not limited to the civilian trauma population. Among combat casualties, sepsis has a prevalence of 30% and lethality of 47% despite aggressive interventions (4). These statistics underscore the urgent need for more effective therapies for trauma-associated sepsis.

Severe trauma and sepsis elicit a systemic inflammatory response, leading to endothelial injury, coagulopathy and loss of vasomotor tone (5–7). A primary marker of endothelial injury in trauma and sepsis is the loss of the endothelial glycocalyx, a gel-like layer lining the blood vessels (8, 9). The glycocalyx is formed by membrane-bound sulfated proteoglycans, consisting of a core protein, syndecan-1, with glycosaminoglycan side chains (8, 9). Syndecan-1 functions primarily as the acceptor for heparan sulfate-binding molecules such as fibrinogen to form the protective glycocalyx (9–11). In severe trauma and sepsis, the extracellular domain of syndecan-1 is cleaved and released into the bloodstream, where the soluble molecule plays a role in modulating systemic inflammation (8, 12). Syndecan-1 shedding is mediated by “sheddases,” among which metalloproteinase 9 (MMP9) is a key mediator after both trauma and sepsis (8, 12–14).

With the advent of balanced blood product-based resuscitation for hemorrhagic shock, which entails the early use of fresh frozen plasma (FFP) in an equal ratio to red cells, came the observation of improved outcomes and reduced organ dysfunction (15–17). In animal models of hemorrhagic shock, the benefits of plasma were associated with restoration of the glycocalyx and reduced syndecan-1 shedding (18, 19). Further, we have previously demonstrated that fibrinogen, a key plasma component, is protective on the endothelium by stabilizing syndecan-1 on the cell surface and maintaining vascular integrity (11, 13).

Therefore, the objective of this study was to investigate the protective effects of fresh frozen plasma (FFP) and fibrinogen (FIB) in a mouse "double-hit" model of combined polymicrobial abdominal sepsis and hemorrhagic shock. We hypothesized that both fibrinogen- and plasma-based resuscitation would mitigate the detrimental effects on lung function through restoration of cell surface syndecan-1.

## Materials and methods

### Animal model of CLP and hemorrhage shock

C57BL/6J mice were obtained from Jackson Laboratory (Bar Harbor, ME). Both male and female adult mice were used at an age of 9 -10 weeks with a body weight of 25-30 g. Mice underwent isoflurane anesthesia via face mask maintaining a dose of 1.5%. Both male and female adult C57BL/6 mice were used. Mice underwent isoflurane anesthesia. The femoral artery and vein were cannulated for continuous blood pressure monitoring and blood withdrawal or fluid administration. Next, a laparotomy was performed for cecal ligation and puncture (CLP). The cecum was exposed and ligated 3mm from its distal tip. The ligated portion was punctured with a 25-gauge needle, and the cecum was returned to the abdomen, which was then closed.

Following CLP, mice underwent hemorrhagic shock (HS) as we described previously (13). Sham mice underwent anesthesia and vessel cannulation but no surgical procedures or fluid administration. The mouse experiments were performed randomly for Sham, Lactated Ringer’s (LR), FIB, and FFP groups. Fibrinogen and FFP were prepared as we described previously (13) and were obtained from CSL Behring GmbH (RioSTAP, Germany) and Tennessee Blood Services (Memphis, TN), respectively. At 24 h after CLP+HS, mice underwent isoflurane anesthesia for vessel cannulation for measurement of mean artery pressure (MAP) and were then euthanized by exsanguination. The blood and lungs were harvested for assays.

### Lung alveolar thickness

Lung tissues were sectioned by a cryostat and stained with hematoxylin and eosin (H&E). Two individuals scored alveolar wall thickness blinded to group assignment, in which 0 indicated normal and 3 indicated maximal thickening.

### Lung tissue immunostaining

To detect lung syndecan-1, myeloperoxidase (MPO), neutrophil elastase, and MMP9, lung tissue cryostat sections were stained with anti-mouse syndecan-1 (sc12765, Santa Cruz Biotechnology), anti-mouse MPO antibody (sc390109, Santa Cruz Biotechnology), anti-mouse neutrophil elastase (sc-55549, Santa Cruz Biotechnology), anti-mouse MMP9 (sc393859, Santa Cruz Biotechnology, which can detect both pro-MMP9 and MMP9) and Alexa Fluor 488 goat anti-mouse IgG (A21121, Invitrogen). Random image acquisition and fluorescence intensity quantification were performed as previously described (13).

### Plasma parameters

Plasma concentrations of syndecan-1 (EK1339, Boster), MPO (MBS2701466, MyBioSource), neutrophil elastase (EK1445, Boster), and MMP9 (EK0466, Boster) were evaluated with respective ELISA kits per manufacturers’ instructions.

### Statistical analysis

Statistical analysis was conducted as we described previously (13). The data were expressed as means ± SE and were analyzed using ANOVA with Tukey’s post-hoc multiple comparison test. P < 0.05 was considered statistically significant.

## Results

### FFP, but not fibrinogen, decreased syndecan-1 shedding

Increased plasma syndecan-1 is an early marker of endothelial injury (19, 20). Our results indicated that 24 hours after the onset of CLP+HS with LR resuscitation, plasma syndecan-1 levels were significantly increased compared to shams (Fig. 1A). Resuscitation with fibrinogen did not alter shedding, whereas resuscitation with FFP decreased plasma syndecan-1 to sham levels (Fig. 1A). Conversely, CLP+HS with LR significantly decreased lung tissue syndecan-1 immunofluorescent intensity compared to shams (Fig 1b and 1C). Resuscitation with fibrinogen had no effect, whereas resuscitation with FFP significantly increased lung syndecan-1 staining compared to LR-treatment (Fig. 1B and 1C).

**Fig. 1.**
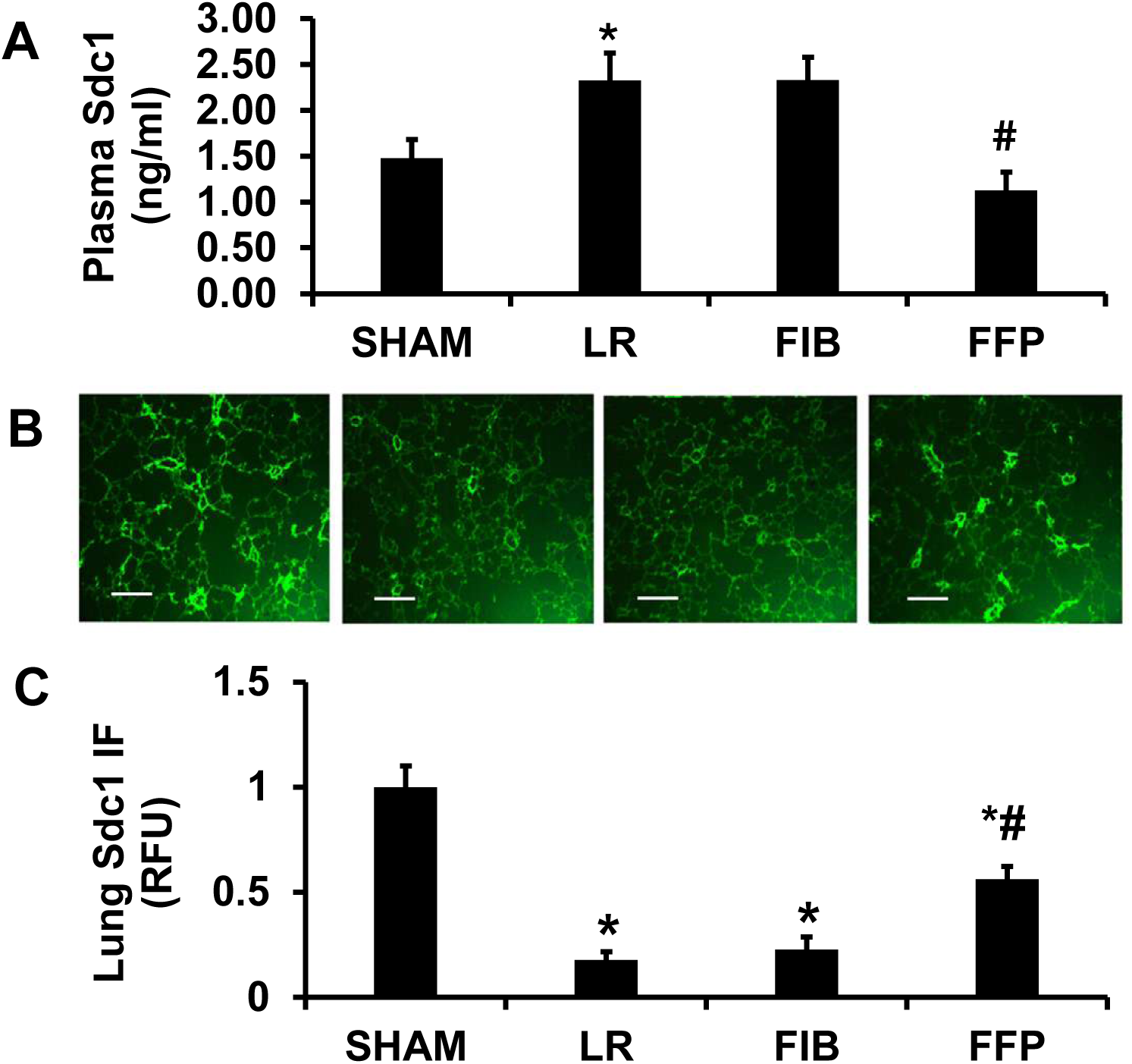
FFP, but not FIB, blocked CLP+HS-induced sydecan-1 shedding. A. Plasma syndecan-1 levels in mice at 24h post CLP+HS. B. Representative lung tissue syndecan-1 immunofluorescent staining (scale bar: 200 µM). C. Quantification of lung tissue syndecan-1 immunofluorescent intensity. N = 8 mice/group; mean ± SEM; ANOVA with post hac Tukey’s test. LR: CLP+HS+lactated Ringer’s solution; FIB: CLP+HS+fibrinogen; FFP: CLP+HS+fresh frozen plasma. * Compared with SHAM, p<0.05; compared with LR, p<0.05.

### FFP, but not fibrinogen, lessened CLP+HS-induced lung injury

Lung alveolar wall thickness is a measurement of lung injury and endothelial permeability. A trend of increased lung alveolar wall thickness was observed after CLP+HS and resuscitation with either LR or fibrinogen compared to shams (Fig. 2A and 2B). However, this increase was significantly attenuated by resuscitation with FFP (Fig. 2A and 2B).

**Fig. 2.**
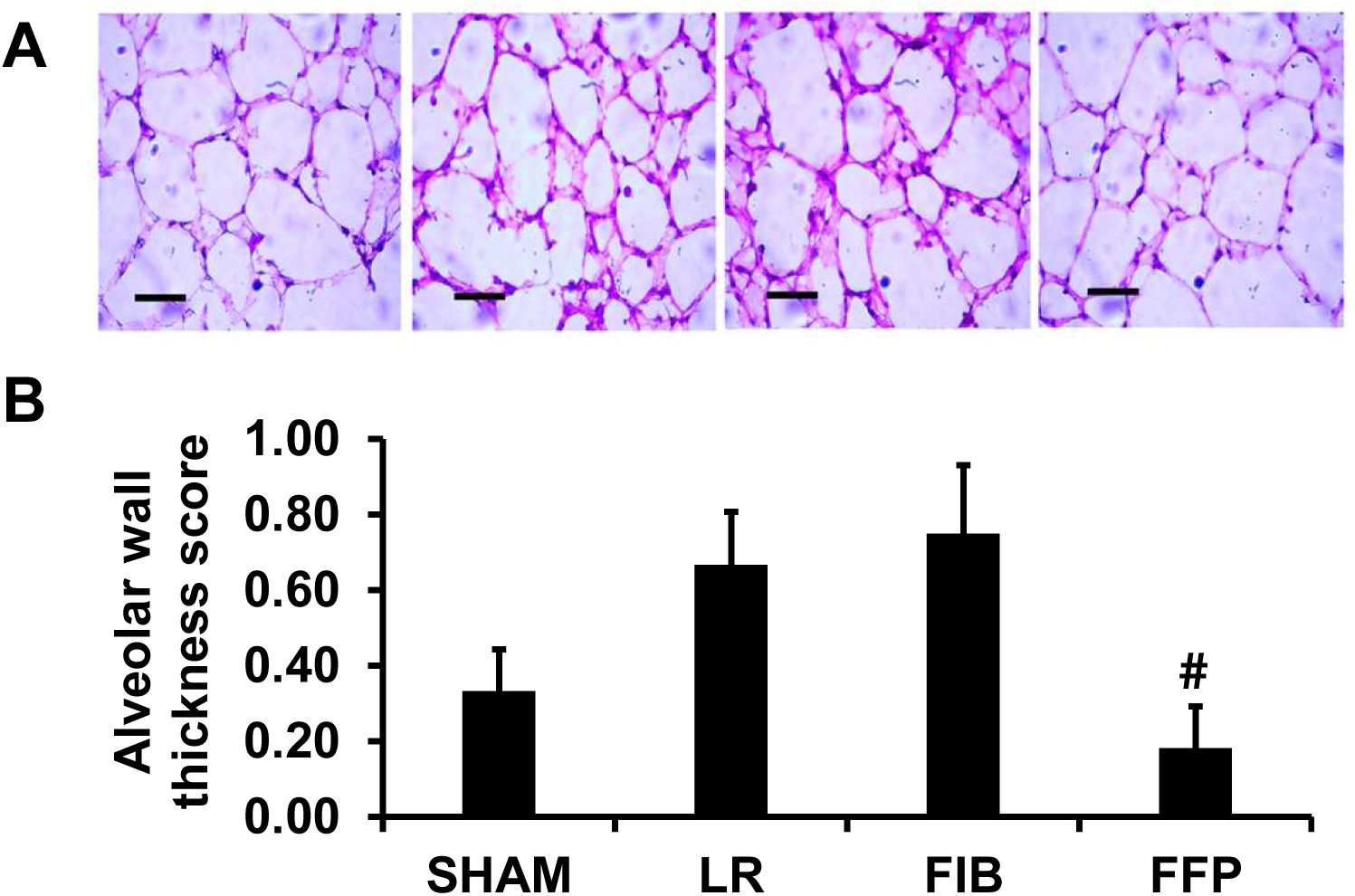
FFP, but not FIB, blocked CLP+HS-induced lung alveolar wall thickness. A. Representative lung section H & E staining in mice at 24h post CLP+HS (scale bar: 50 µM). B. Lung alveolar wall thickness scores. N = 8 mice/group; mean ± SEM; ANOVA with post hac Tukey’s test. # compared with LR, p<0.05.

### CLP+HS induced lung neutrophil degranulation, which was partially blocked by FFP but not fibrinogen

MPO, neutrophil elastase, and pro-MMP9 are enzymes stored in the granules of neutrophils and monocytes and released into the circulation upon degranulation (21–23). Once released systemically, neutrophil elastase cleaves the pro-domain (the inhibitory section) of pro-MMP9, which causes a structural change to expose its catalytic site such that the active MMP9 state exists in circulation (21–23). MMP9 is a known sheddase for syndecan-1 during hemorrhagic shock and sepsis (13, 14). Compared with shams, CLP+HS with LR significantly increased plasma levels of MPO (Fig. 3A), neutrophil elastase (Fig. 4A) and MMP9 (Fig. 5A). These parameters were all significantly reversed by resuscitation with FFP, but not fibrinogen.

**Fig. 3.**
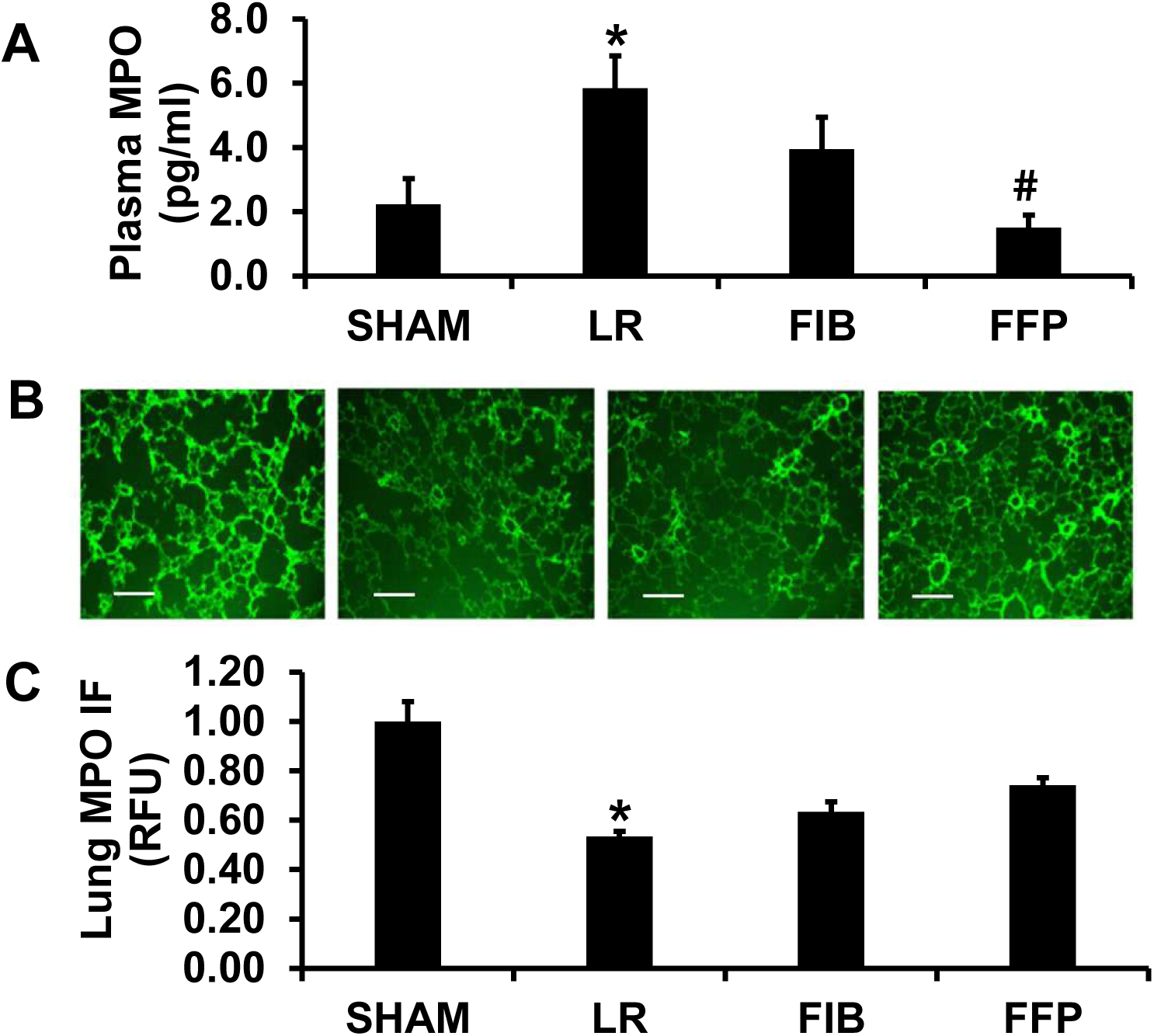
FFP, but not FIB, blocked CLP+HS-induced lung neutrophil MPO degranulation. A. Plasma MPO levels in mice at 24h post CLP+HS. B. Representative lung tissue MPO immunofluorescent staining (scale bar: 200 µM). C. Quantification of lung tissue MPO immunofluorescent intensity. N = 8 mice/group; mean ± SEM; ANOVA with post hac Tukey’s test. * Compared with SHAM, p<0.05; compared with LR, p<0.05.

**Fig. 4.**
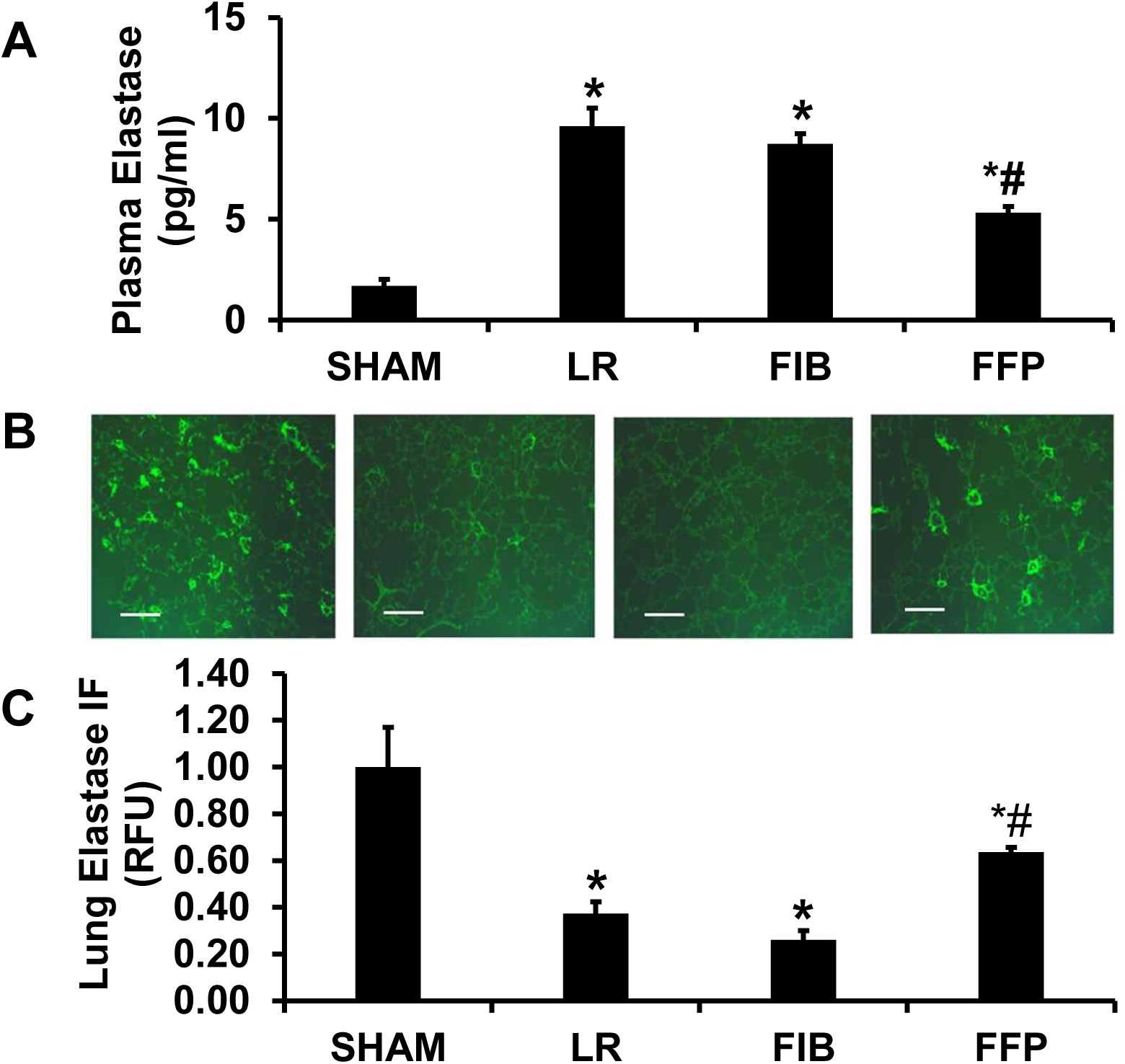
FFP, but not FIB, blocked CLP+HS-induced lung neutrophil elastase degranulation. A. Plasma neutrophil elastase levels in mice at 24h post CLP+HS. B. Representative lung tissue elastase immunofluorescent staining (scale bar: 200 µM). C. Quantification of lung tissue elastase immunofluorescent intensity. N = 8 mice/group; mean ± SEM; ANOVA with post hac Tukey’s test. * Compared with SHAM, p<0.05; compared with LR, p<0.05.

**Fig. 5.**
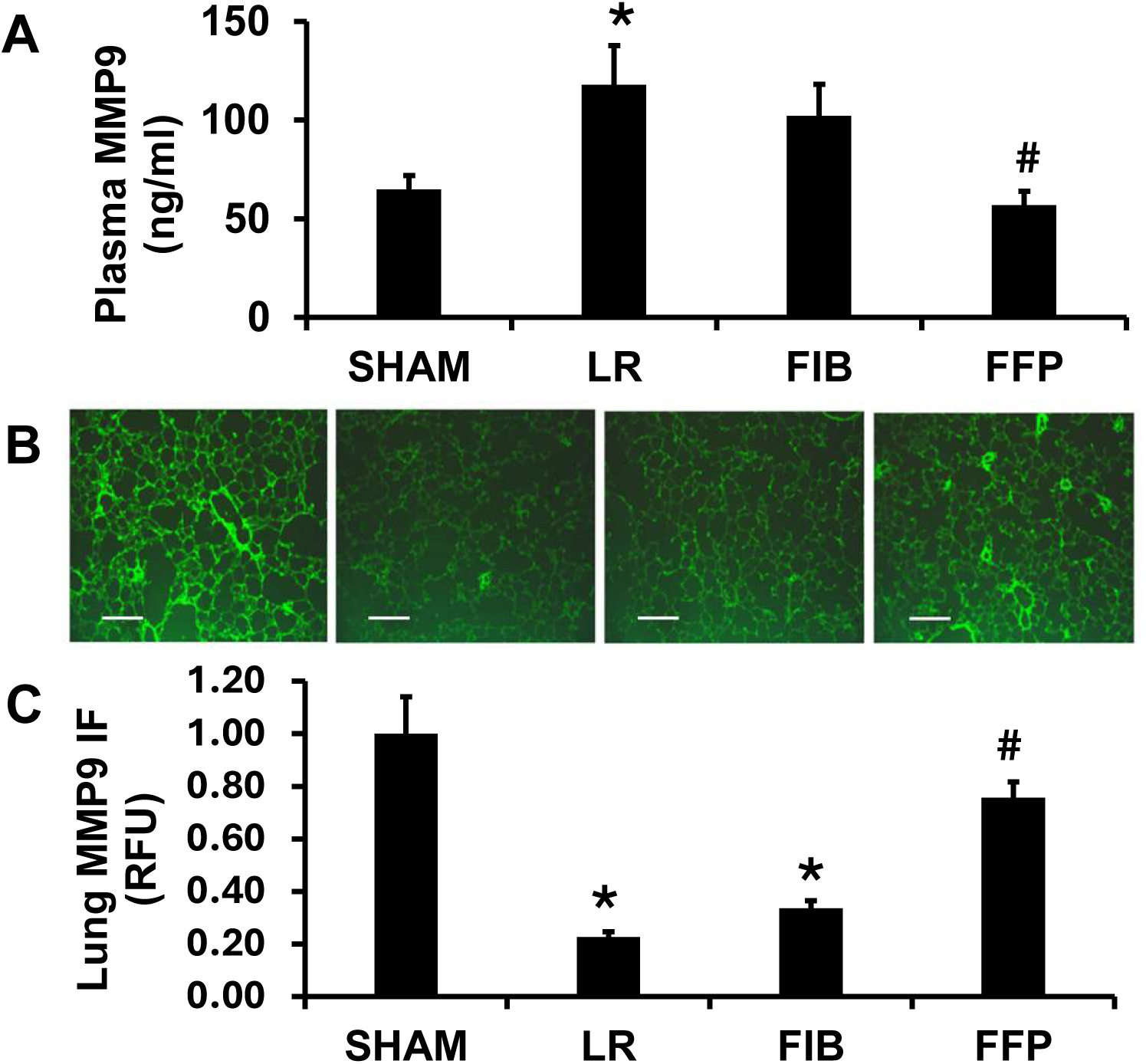
FFP, but not FIB, blocked CLP+HS-induced lung neutrophil MMP9 degranulation. A. Plasma MPP9 levels in mice at 24h post CLP+HS. B. Representative lung tissue MMP9 immunofluorescent staining (scale bar: 200 µM). C. Quantification of lung tissue MMP9 immunofluorescent intensity. N = 8 mice/group; mean ± SEM; ANOVA with post hac Tukey’s test. * Compared with SHAM, p<0.05; compared with LR, p<0.05.

Consistently, lung immunofluorescent staining indicated that, compared with sham, CLP+HS with LR significantly decreased lung tissue MPO (Fig. 3A and 3B), lung tissue neutrophil elastase (Fig. 4A and 4B), and lung tissue MMP9 (Fig. 5A and 5B). Importantly, these decreases were again partially reversed by FFP, but not fibrinogen. These data indicate that CLP+HS caused reciprocal changes in the levels of MPO, neutrophil elastase, and MMP9 between the lung tissue and plasma. Further, the results suggest that FFP acts through inhibiting CLP+HS-induced degranulation by neutrophils and monocytes.

### FFP, but not fibrinogen, partially reversed CLP+HS-induced hypotension

The above data indicate that FFP protects the endothelium by maintaining endothelial syndecan- 1/glycocalyx and endothelial barrier, which is known to play a critical role in modulating blood pressure (24). We thus measured the blood pressure in mice 24 hours after CLP+HS and resuscitation with LR (vehicle control), fibrinogen or FFP. CLP+HS significantly decreased MAP compared with shams (97 ± 2.7 mmHg Sham vs 40 ± 6.7 mmHg LR*; n=8, *p<0.05 compared with Sham), indicating that the combined abdominal sepsis and hemorrhagic shock model resulted in hypotension. CLP+HS-induced hypotension was significantly improved by FFP but not by fibrinogen (40 ± 6.7 mmHg LR vs 53 ± 5.3 mmHg fibrinogen vs 76 ± 4.5 mmHg FFP^#^; n=8, ^#^p<0.05 compared with LR). Summary of CLP+HS results is shown on Fig. 6.

**Fig. 6.**
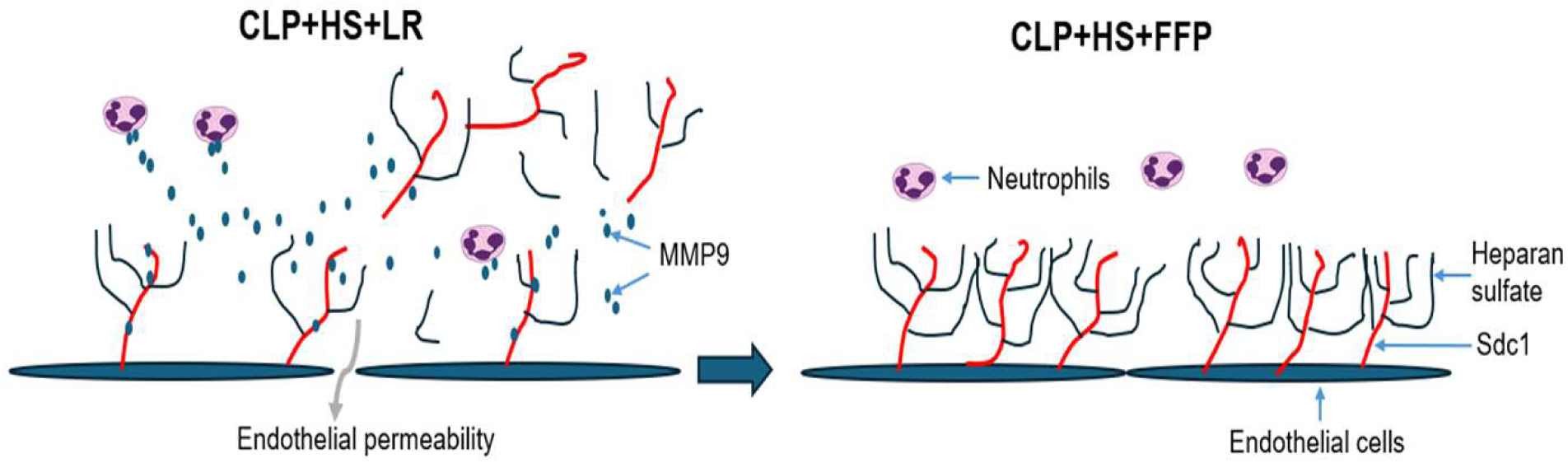
Summary of the results. CLP+HS caused global neutrophil degranulation, releasing MMP9 to cleave endothelial syndecan-1, which was blocked by FFP.

## Discussion

The current study demonstrated that a combined model of polymicrobial abdominal sepsis and hemorrhagic shock resulted in lung injury, as demonstrated by an increase in lung histopathologic injury, and decreases in lung tissue syndecan-1, MPO, neutrophil elastase, and MMP9, with reciprocal elevations in plasma syndecan-1, MPO, neutrophil elastase, and MMP9. These detrimental alterations were significantly attenuated by FFP but not by fibrinogen.

Polymicrobial sepsis and hemorrhage shock both trigger proinflammatory cytokine production that activate neutrophils and monocytes to release proteases (e.g., neutrophil elastase and MMP9) through degranulation (25). Though these proteases kill bacteria in the case of sepsis, they may also result in tissue injury. For example, the released neutrophil elastase activates MMP9 in the circulation (21, 26), which in turn cleaves syndecan-1/glycocalyx and results in endothelial injury to targeted organs such as the lung (12, 14).

Although plasma has clearly been shown to benefit the endothelium after trauma and hemorrhagic shock, preclinical and clinical studies in sepsis are not definitive. In a rat model of CLP sepsis, Chang et al. found that plasma increased 48-hour survival, improved pulmonary function, and reduced pulmonary edema compared to saline (27). In a similar mouse model of cecal slurry-induced abdominal sepsis, we were able to only demonstrate decreases in pro-inflammatory gene expression but not in the indices of pulmonary dysfunction (28). There is only one small single center clinical study that examined the use of a pathogen inactivated plasma product in patients with septic shock (29). Plasma was found to be safe and feasible, but the authors were unable to demonstrate any improvement in microvascular perfusion or in the biomarkers of endothelial activation/dysfunction, for which the small size and heterogeneity of the study may have contributed (29).

Previously we showed that fibrinogen but not plasma was protective on endothelial syndecan-1 and lung permeability in a combined model of pneumonia and hemorrhagic shock (13).

However, in the current study, we found that only plasma afforded protection. In our pneumonia and hemorrhagic shock model, mice were in a state of innate immunity deficiency as indicated by a decrease in lung neutrophils which are needed for bacterial clearance. Thus, fibrinogen supplementation was able to maintain lung neutrophils and play a role in host defense (13).

Moreover, fibrinogen protected lung endothelial syndecan-1/glycocalyx from cleavage and reduced lung permeability (13). In contrast, the current combined model of abdominal sepsis and hemorrhage shock triggered a systemic proinflammatory response that can drive robust reactive oxygen species production via neutrophil and monocyte respiratory burst (30). As circulating fibrinogen is highly prone to oxidation (31–33), the administered fibrinogen was likely rapidly oxidized and crosslinked, leading to a loss of its protective function.

The protective mechanism of FFP may involve restoration of plasma degranulation inhibitors, such as histidine-rich glycoprotein (HRG) and inter-α-inhibitor proteins (IAIPs) (34), which are depleted during hemorrhagic shock and sepsis. Additionally, plasma components like protease inhibitors (e.g., alpha-1 antitrypsin) can directly bind and neutralize neutrophil granule-derived proteases such as neutrophil elastase (35). This is significant because neutrophil elastase plays an important role in converting pro-MMP9 to its active form, MMP9 (21, 26). By reconstituting the protective plasma proteome, FFP may inhibit neutrophil degranulation, limit the release of cytotoxic granular enzymes, and mitigate endothelial injury, thereby contributing to its therapeutic benefit.

There are several limitations in the current study. In contrast to our previous observations in pneumonia and HS (13), fibrinogen did not protect the lung endothelium in the present CLP+HS model. The precise reason for this discrepancy remains to be determined. Second, we only examined changes at 24 hours after CLP+HS rather than at serial timepoints. Lastly, we only evaluated the effects of FFP on the lungs, as they are the most frequently injured organs after trauma, while the kidney and heart are more likely to be affected by sepsis (36, 37).

In summary, in the current CLP+HS model, FFP, but not fibrinogen, conferred significant endothelial and pulmonary protection. FFP resuscitation restored hemodynamic stability, attenuated lung injury and endothelial permeability, and preserved glycocalyx integrity as evidenced by normalization of syndecan-1 levels in both plasma and lung tissue.

Mechanistically, CLP+HS induces global neutrophil degranulation to release elastase, which then mediates the conversion of pro-MMP9 to its active form and thereby causes lung endothelial syndecan-1 shedding and lung injury. FFP resuscitation suppresses neutrophil and monocyte degranulation, with consequent reduction in circulating neutrophil elastases, myeloperoxidase, and active MMP9. These findings support the therapeutic use of fresh frozen plasma following traumatic hemorrhagic shock with sepsis.

## Funding declaration

This work was funded by the US Air Force grant 30039991.

The views and opinions presented herein are those of the author(s) and do not necessarily represent the views of DoD or its Components. Unless otherwise noted, imagery in this document is property of the U.S. Air Force.

## Author Contributions

FW and RK conceptualized the study, wrote the proposal, and obtained ethical approval. FW and CR performed the experiments and collected the data. FW, RK and JC analyzed data and wrote the manuscript. All authors reviewed and approved the final manuscript.

## Data Availability

The datasets generated and/or analyzed during this study are available from the corresponding author upon reasonable request.

## Animals use disclaimer

Research was conducted in accordance with an approved animal use protocol at an AAALAC International-accredited facility, in full compliance with all applicable federal statutes and regulations governing the use of animals in research. The study adhered to the principles outlined in the Guide for the Care and Use of Laboratory Animals (National Research Council, 2011 edition). The Animal Care and Use Committee of Institutes of the University of Maryland School of Medicine approved the animal use protocol (IACUC #0422005).

## Human blood components use disclaimer

The Institutional Review Board approved the use of human blood components in mice (IBC #00006292). The board contact: John O’Neill. There was no direct involvement of human participants in the study.

